# Adenine base editing is an efficient approach to restore function in FA patient cells without double-stranded DNA breaks

**DOI:** 10.1101/2022.04.22.489197

**Authors:** Sebastian M. Siegner, Alexandra Clemens, Laura Ugalde, Laura Garcia-Garcia, Juan A. Bueren, Paula Rio, Mehmet E. Karasu, Jacob E. Corn

**Affiliations:** Department of Biology, ETH Zurich, Zurich 8053, Switzerland; Division of Hematopoietic Innovative Therapies, Centro de Investigaciones Energéticas Medioambientales y Tecnológicas (CIEMAT), Madrid 28040, Spain

**Author notes:** These authors contributed equally (Sebastian M. Siegner, Alexandra Clemens, Laura Ugalde).

## Abstract

Fanconi Anemia (FA) is a debilitating genetic disorder with a wide range of severe symptoms including bone marrow failure and predisposition to cancer. CRISPR-Cas genome editing manipulates genotypes by harnessing DNA repair and has been proposed as a potential cure for FA. But FA is caused deficiencies in DNA repair itself, preventing the use of editing strategies such as homology directed repair. Recently developed base editing (BE) systems do not rely on double stranded DNA breaks and might be used to target mutations in FA genes, but this remains to be tested. Here we develop a proof of concept therapeutic base editing strategy to address two of the most prevalent FANCA mutations in patient cells. We find that optimizing adenine base editor construct, vector type, guide RNA format, and delivery conditions lead to very effective genetic modification in multiple FA patient backgrounds. Optimized base editing restored FANCA expression, molecular function of the FA pathway, and phenotypic resistance to crosslinking agents. ABE8e mediated editing in primary hematopoietic stem and progenitor cells from an FA patient was both genotypically effective and restored FA pathway function, indicating the potential of base editing strategies for future clinical application in FA.

## Introduction

Fanconi Anemia (FA) is a serious genetic disorder mainly characterized by developmental abnormalities, cancer predisposition and bone marrow failure (BMF), which becomes evident in most FA patients during the first decade of life ^1,2, 3^. Androgen therapy and regular blood transfusions can ameliorate the BMF in FA patients, but do not address the underlying cause of the disease ^4^. Allogeneic hematopoietic stem cell transplantation (HSCT) from healthy donors is the only curative treatment of BMF in these patients, but carries serious risks such as graft-vs-host disease and increased incidence of squamous cell carcinoma in the long term ^5, 6^. Genetic treatments to complement or repair the mutations that cause FA during autologous HSCT would offer many benefits over traditional therapy, potentially providing a lasting cure without the side effects associated with allogeneic BMT ^7^

FA is caused by mutations in any of the 23 genes that participate in the FA/BRCA DNA damage response pathway ^8^. The FA gene products work together in physical complexes and connected pathways to repair interstrand cross links (ICLs) in DNA, which can be caused by DNA damaging agents such as chemotherapeutics (cisplatin, mitomycin C) or endogenous metabolic byproducts such as aldehydes ^9, 10^. In the absence of a functional FA pathway, these unresolved ICLs eventually lead to chromosomal breaks and genome instability. FA patient cells are also compromised by normal levels of pervasive stressors such as replication fork collapse ^11^, emphasizing the importance of the FA pathway to guardian genome integrity.

Lentiviral mediated gene therapy has recently been successfully used to treat HSCs from *FANCA-*deficient (FA-A) patients, the most prevalent FA complementation group, with the help of optimized HSC mobilization ^12^ and transduction protocols ^13^. Although no severe side effects have been observed neither in the FA gene therapy trial, or in most of the LV-mediated gene therapy trial developed so far ^14^, the possibility of conducting precise gene repair in mutated sequences accounting for a genetic disease, in FA HSCs in this particular case, offers an additional safeguard to improve the safety of gene therapy. Furthermore, gene editing would also maintain endogenous regulation of gene expression and could extend therapeutic application to FA complementation groups associated with mutations in strongly regulated or genes.

“Classic” CRISPR-Cas genome editing relies on creating a targeted DNA double-strand break (DSB) that can be resolved by either error-prone pathways to create semi-random small insertions and deletions (indels), or by precise homology directed repair (HDR) from a template ^15, 16^. Although HDR could theoretically facilitate to “surgically” change almost any mutation to the wild type sequence, its efficiency is low in primitive HSCs and is particularly compromised in FA-HSCs, due to their defects in HDR ^17^. Indel-based genome editing has been demonstrated to be a good alternative to correct specific FA mutations, converting nonsense to in-frame mutations that restore the FA gene function ^18^. But this approach is limited in the spectrum of patient mutations that could be addressed.

Newer genome editing systems such as base editing (BE) that work without inducing double stranded DNA breaks theoretically offer great opportunities to precisely correct specific mutations in the FA genes ^19, 20^. Nevertheless, whether a path to a genetic cure for FA is feasible while using one of the many existing base editors is unclear. Here we report for the first time a BE approach to address two of the most prevalent *FANCA* mutations in patient cells. We found that adenine base editing can be remarkably effective to target FA alleles. Optimization of the adenine base editor construct, vector type, guide RNA format, and delivery conditions restored FANCA expression, molecular function of the FA pathway, and phenotypic resistance to crosslinking agents. Importantly, ABE8e induced unprecedentedly high levels of gene conversion in HSPCs from a FA patient, confirming the great potential of this strategy for the future clinical application in FA.

## Results

To develop a proof-of-concept base editing therapy for FA, we first employed lymphoblastoid cell lines (LCLs) generated from either healthy donors (HD) or FA patients (Figure 1A). These immortalized cells recapitulate the major phenotypic hallmarks of FA, including reduced proliferation and sensitivity to crosslinking agents, and allowed us to test the efficacy and toxicity of different tools and protocols of BE. Mutations in *FANCA* account for approximately 60-65% of FA ^21^, and we focused on two prevalent alleles of *FANCA* ^22, 23^. FA-55 is an FA LCL carrying a homozygous mutation in *FANCA* gene which consists on a premature stop codon at exon 4 (c.295C>T). On the other hand, FA-75 harbors an compound heterozygous mutation in *FANCA* gene (c.2639G>A and c.3788_3790 del TCT)^18^.

**Figure 1:**
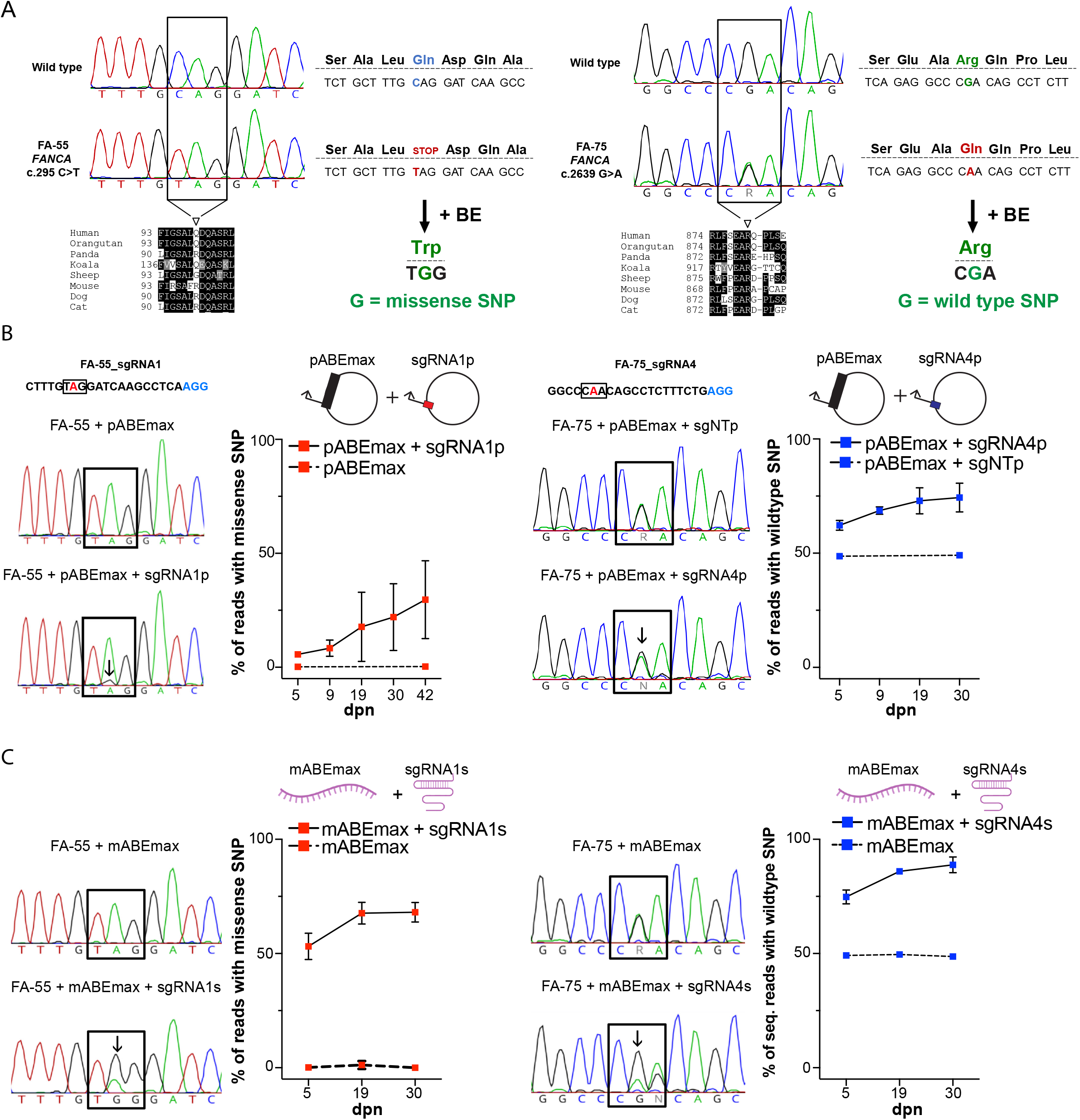
Base editing is an efficient approach to modify Fanconi Anemia mutations. **A)** Details of FA-55 and FA-75 mutations. Sanger traces from wild type and mutant LCLs showed the indicated c.295 C>T and c.2639 G>A mutations. Next to Sanger traces, translation of codons is illustrated for each mutation. FA-55 c.295 C>T mutation leads to stop codon, terminating translation of FANCA prematurely. Adenine base editing is designed to introduce a missense SNP that encodes a Tryptophan instead of a Glycine. FA-75 c.2639 G>A leads to Arginine to Glycine mutation. Adenine base editing reverts the missense SNP to wild type sequence. FA-75 is compound heterozygous and the wild type SNP is already present in unedited cells. Below Sanger traces, protein sequence alignments from multiple species are shown. Amino acids with dark background or grey background indicate identity or similarity among different species, respectively. **A)** Base editing in FA-55 and FA-75 LCLs by delivering ABEmax and sgRNA in plasmid format. On the left top side, the FA-55 sgRNA1 target site is shown. PAM sequence and edited base are highlighted by blue and red fonts, respectively. The FA-75 sgRNA4 site is shown. Representative Sanger traces show initial editing 5 days after electroporation of FA-55 and FA-75 LCLs, all with ABEmax and with or without the indicated sgRNAs. In the case of FA-75 targeting, plasmid carrying non targeting sgRNA (sgNTp) was included. Arrows indicate presence of edited alleles. Graphs show edited allele frequency measured by amplicon NGS in a time course after editing. Continuous lines represent the pool of cells electroporated with base editor and sgRNA, while dashed lines represent the pool of cells electroporated with base editor alone. The graphs summarize 2 biological replicates, error bars indicate the range of values. **B)** Base editing of FA-55 and FA-75 LCLs by delivering ABEmax and sgRNA in mRNA format. Representative Sanger traces show initial editing 5 days after electroporation with ABEmax with or without sgRNAs. Arrows indicate presence of edited alleles. Graphs show edited allele frequency measured by amplicon NGS in a time course after editing. Continuous lines represent the pool of cells electroporated with base editor and sgRNA, while dashed lines represent the pool of cells electroporated with base editor alone. The graphs summarize 2 biological replicates, error bars indicate the range of values.

Both the FA-55 and FA-75 mutations are not amenable to cytosine base editing (CBE) due to the identity and context of the mutations, but could theoretically be addressed with adenine base editing (ABE)(Figure 1A). The FA-75 G-to-A mutation could be reverted back to wildtype by targeting the adenine mutation in the coding strand. The FA-55 T-to-C mutation might also be reverted to wildtype by ABE targeting on the non-coding strand. But this mutation is very close to several other non-wobble coding strand thymidines that lie in the base editing window (Figure 1A). Modification of these positions would lead to coding changes expected to impair protein function. Therefore, we focused on editing the coding strand, in which fewer potentially deleterious bystander mutations could occur. Our strategy aimed to convert the FA-55 nonsense mutation to a tryptophan missense mutation. The targeted amino acid of *FANCA* is particularly non-conserved (Figure 1A) and expendable for FANCA function ^18^, so we reasoned that this change might be tolerated for FANCA function. We used PnB Designer to design several candidate gRNAs to base edit each FA genotype^24^.

Base editors have rapidly diversified and multiple next-generation ABEs are available. ABEmax is a second-generation adenine base editor that has been used in several contexts ^25^ and has been characterized extensively to establish its targeting window and off-target propensities ^26–28^. We tested whether delivery of candidate gRNAs and ABEmax in plasmid format yielded intended base editing in FA-55 and FA-75 contexts. Bulk Sanger sequencing and next generation Illumina sequencing of PCR amplicons five days after electroporation of FA-55 and FA-75 LCLs indicated low levels of base editing (5.66 ± 0.59% A to G) for FA-55. In the case of FA-75, a 62.20 ± 2.12% A to G conversion was observed. Nevertheless, FA-75 harbors a heterozygous mutation and so has a baseline wildtype level of 50% at the targeted sites (Figure 1B, Sanger traces).These results suggested that these guide and base editor combinations were capable of genomic targeting to induce the desired sequence changes, though with modest efficiency in the current format.

To determine whether the respective edited alleles conferred proliferative advantage over cells harboring the mutant alleles, we monitored the growth of edited and unedited cells in bulk cultures using next-generation Illumina sequencing of PCR amplicons (amplicon-seq) that target each edited site. During 30 days of culture after editing, conversion of the FA-75 nonsense mutation to wildtype led to increased levels of the wild type base to 74.35 ± 6.35%. Significantly, conversion of the FA-55 nonsense mutation to missense also led to a proliferative advantage for edited cells, increasing to 29.67 ± 17.09% of the altered base. (Figure 1B, solid lines). Edited reads were not found in cells kept in culture for the same length of time but electroporated with only base editor and no gRNA, indicating that spontaneous reversion did not play a role in outgrowth of cells with the wildtype sequence. (Figure 1B, dashed lines)

Since double stranded DNA plasmid delivery is inefficient and highly toxic in HSCs ^29^, we next tested electroporation of an mRNA coding for ABEmax together with a chemically protected, synthetic gRNA. This combination has been effective in editing HSPCs from non-FA genotypes ^30–32^.

We found that mRNA ABEmax and synthetic gRNA dramatically increased bulk editing levels soon after electroporation for FA-55, with the mutation now contributing to the majority of the Sanger sequencing chromatogram (Figure 1C) and no qualitative evidence of bystander mutations. mRNA based editing of the FA-75 allele was also improved relative to plasmid editing (Figure 1C, right, bottom Sanger tracks). However, FA-75 editing was associated with a bystander mutation at the adjacent 3’ adenine in the wobble position. We further quantified editing efficiency at each locus using amplicon-seq. mRNA delivery of ABEmax paired with synthetic guide RNAs yielded high levels of editing in both FA-55 (missense 53.14 ± 5.77% desired base and FA-75 correction, 74.75 ± 3.04% desired base) after 5 days in culture (Figure 1C). In longer-term cultures, edited allele frequencies steadily increased for both *FANCA* genotypes, representing the great majority of reads after 30 days. Taken together, these results indicate that using mRNA delivery of ABEmax paired with a synthetic guide RNA can be very effective at genetic modification in two different *FANCA* genotypes. Our results also suggest that the *FANCA* missense edit we tested here is capable of providing a fitness benefit relative to cells with FA-55 c.295C>T mutation.

We next asked if the base edited LCLs have restored FA pathway function (Figure 2A). Cells derived from FA patients exhibit hypersensitivity to DNA interstrand crosslinking reagents such as mitomycin C (MMC) and cisplatin ^33^. Unedited FA-55 and FA-75 LCLs were both hypersensitive to MMC compared to healthy donor LCLs (Figure 2B). To test if gene editing modified the MMC-hypersensitivity of these cells, samples were electroporated with ABEmax mRNA and synthetic guide RNAs, passaged for thirty days in culture, and then assayed for MMC sensitivity. At this point, both FA-55 and FA-75 LCLs exhibited complete phenotypic restoration, as compared to wild type LCLs (Figure 2B).

**Figure 2:**
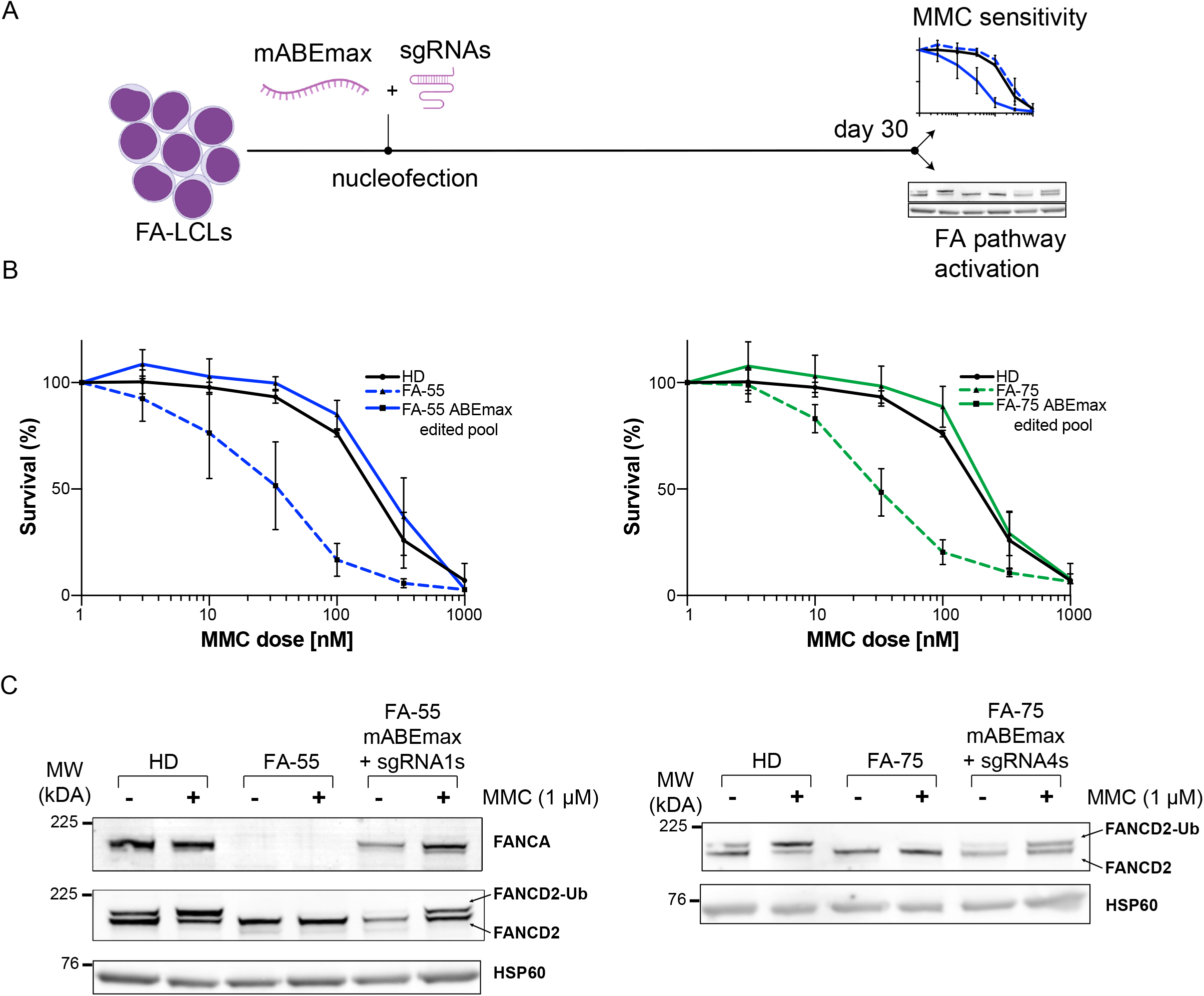
Base editing successfully reverts classical FA phenotypes. **A)** Schematics of experimental design to edit FA-55 or FA-75 LCLs. Cells were edited with base editor mRNA and synthetic gRNA and grown for 30 days in culture. Editing efficiencies were assessed by amplicon NGS, as shown in Figure 1c. **B)** MMC resistance of edited FA-A LCLs. Black line indicates healthy donor (HD) LCL response to increasing doses of MMC. Dashed blue or green lines represent FA-55 or FA-75 LCLs, respectively. Solid blue or green lines represent FA-55 or FA-75 edited pools, respectively. The graphs summarize 3 biological replicates, error bars indicate SD. **C)** Representative Western blots. Indicated cell populations were challenged with 1 μM MMC for 24 hr. Protein extracts were analyzed by Western blot with indicated antibodies (anti-FANCA, anti-FANDC2, anti-HSP60 as a control). FANCD2 or FANCD2-Ub bands are indicated by arrows.

FANCD2 monoubiquitination is a molecular hallmark of FA pathway activation in response to MMC exposure ^34^. In the absence of functional FANCA protein and FA core complex assembly, the FANCD2-FANCI heterodimer cannot be monoubiquitinated. In HD LCLs we verified robust basal expression of FANCA and MMC-induced ubiquitination of FANCD2 (Figure 2C). Neither the FA-55 nor FA-75 LCLs were capable of ubiquitinating FANCD2 in response to MMC treatment. However, bulk ABEmax edited pools of either genotype robustly ubiquitinated FANCD2 after MMC exposure (Figure 2C). Notably, also the missense edit conferred to the FA-55 LCL restored FANCA protein expression and FANCD2 monoubiquitination, further highlighting that the nonsense-to-tryptophan base edit was sufficient to rescue the FA pathway.

While we were in the process of characterizing ABEmax-edited FA LCLs, a hyperactive adenine base editor variant was developed by the labs of Jennifer Doudna and David Liu ^35^. ABE8e was reported to outperform ABEmax in terms of editing efficiency in some cell lines, but with a slightly increased propensity for bystander and off-target effects. We wondered whether ABE8e could further increase base editing levels at early timepoints, especially in FA patient backgrounds, since achieving a very high level of initial editing could be critical when attempting to edit the especially rare HSPCs that can be isolated FA patients.

To compare the efficiencies of ABE8e and ABEmax, we followed a similar experimental design as described in Figure 2. To determine whether the hyperactive ABE8e was more efficient to generate point conversions, without relying on a survival advantage of edited cells, we amplicon-sequenced cell pools only five days after electroporation (Figure 3A). Although even at this short time point, ABE max showed high editing efficiency (28.84 ± 6.95% and 70.43 ± 7.116% in FA-55 and FA-75, respectively), ABE8e further increased this efficiency to 68.44 ± 9.28% and 86.46 ± 3.64%, respectively (Figure 3B and C).

**Figure 3:**
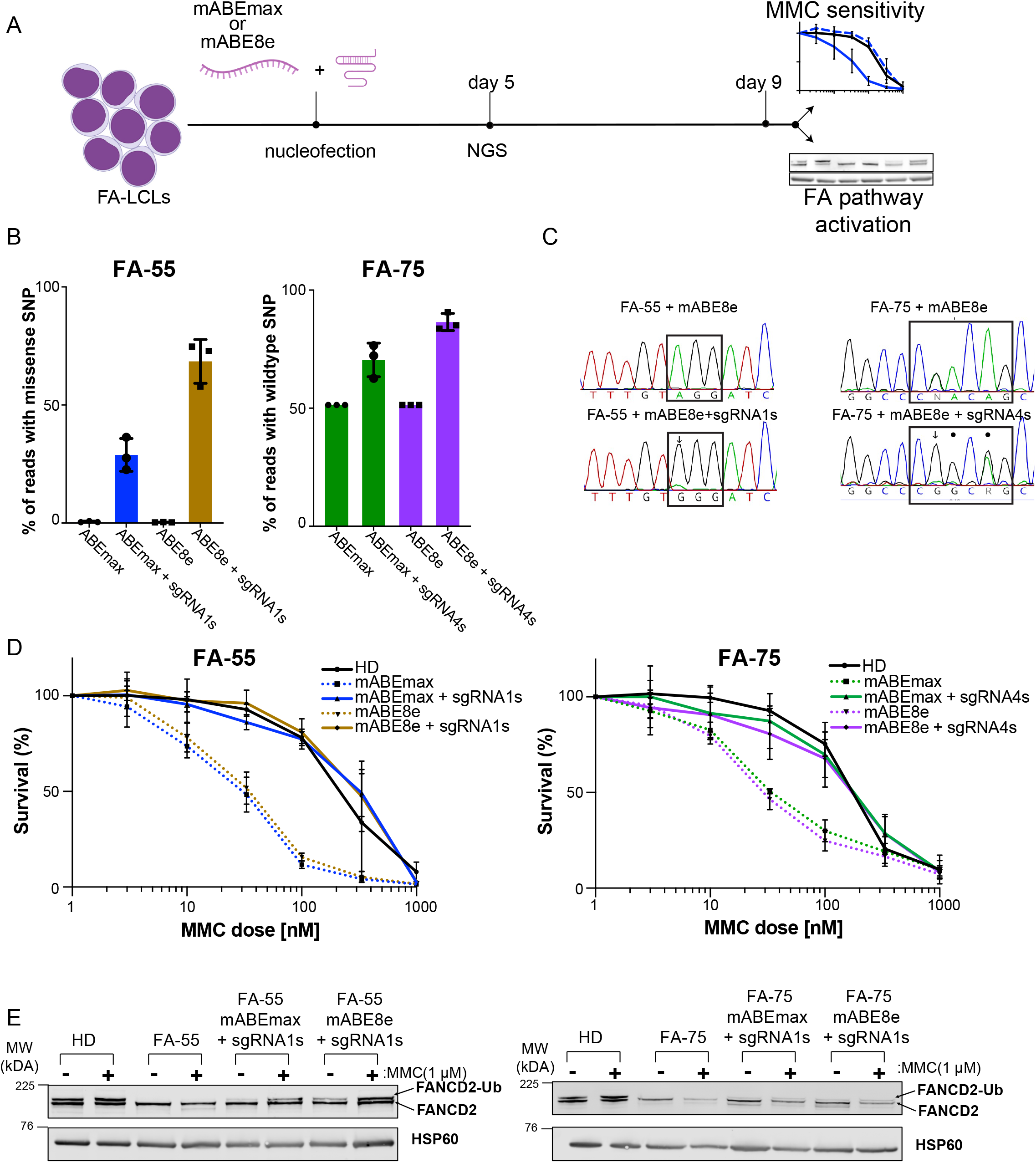
Comparison of ABEmax and ABE8e for editing and phenotypic correction in FA LCLs. **B)** Schematics of experimental design to edit FA-55 or FA-75 LCLs by ABEmax or ABE8e using mRNA base editor and synthetic gRNAs. Electroporated cells were collected at day 5 to measure editing efficiency and at day 9 to measure MMC resistance and activation of FANCD2 ubiquitination. **C)** Quantification of editing levels by amplicon NGS on day 5 in edited LCL populations. Dot or filled squares indicate individual experiments, bars represent the mean of 3 independent experiments, error bars indicate SD. **D)** Representative Sanger traces show initial editing 5 days after electroporation of FA-55 and FA-75 with ABE8e mRNA and with or without indicated synthetic sgRNAs. Arrows indicate presence of edited alleles. Dots indicate presence of bystander edits. **E)** MMC survival of edited FA-A LCLs. Black lines indicate healthy donor (HD) LCL response to increasing doses of MMC. Dashed lines represent FA-55 or FA-75 LCLs electroporated only with ABEmax or ABE8e. Solid colored lines represent FA-55 or FA-75 electroporated with sgRNAs and ABEmax or ABE8e. The graphs summarize 3 biological replicates, error bars indicate SD. **F)** Representative Western blots. Indicated cell populations were challenged with 1 μM MMC for 24 hr. Protein extracts were analyzed by Western blot with indicated antibodies (anti-FANDC2, anti-HSP60). FANCD2 or FANCD2-Ub bands were indicated by the arrows.

Since the correction of a single allele is already sufficient to correct the disease phenotype in FA ^36, 37^, we asked whether ABE8e edited pools also exhibited greater phenotypic correction at short time points when compared to ABEmax edited pools. We thus tested MMC sensitivity and FANCD2 mono-ubiquitination only nine days after editing. (Figure 3A). Nevertheless, editing efficiency reached by either ABE8e and ABEmax resulted in similar levels of MMC resistance, which were equivalent to those observed in HD LCLs (Figure 3D). The phenotypic correction of FA-55 and FA-75 FA LCLs was also supported by the restoration of FANCD2 ubiquitination, in both the ABEmax and ABE8e edited pools (Figure 3E).

Cas-based genome editing tools can affect off-target genomic loci that have sequences similar to the on-target guide RNA ^38^. To further characterize ABE8e and ABEmax editing in FA LCLs, we computationally predicted potential off-target sites for both the FA-55 and FA-75 guide RNAs using Cas-OFF Finder ^39^. For the FA-55-targeting sgRNA1 we found 8 potential sites (Table 1). For the FA-75 targeting sgRNA4 there were 22 potential sites (Table 2). We investigated the top eight potential sites for each guide RNA using amplicon-sequencing. The FA-55 targeting sgRNA1 exhibited no detectable editing in any of the tested candidate off-target sites, irrespective of base editor (Figure 4, top). However, the FA-75 targeting sgRNA4 had a prominent off-target site located on chr2: 202,148,657-202,148,676, with 18.08 ± 1.55% base editing with ABE8e (Figure 4, bottom left graph). ABEmax edited FA-75 cells also had the same off-target site, albeit at lower editing levels 2.79 ± 0.20% (Figure 4, bottom right graph). This site is in the intron 13 of an uncharacterized gene called *KIAA2012*. The potential effects of editing at this off-target remain to be determined.

**Table1:**
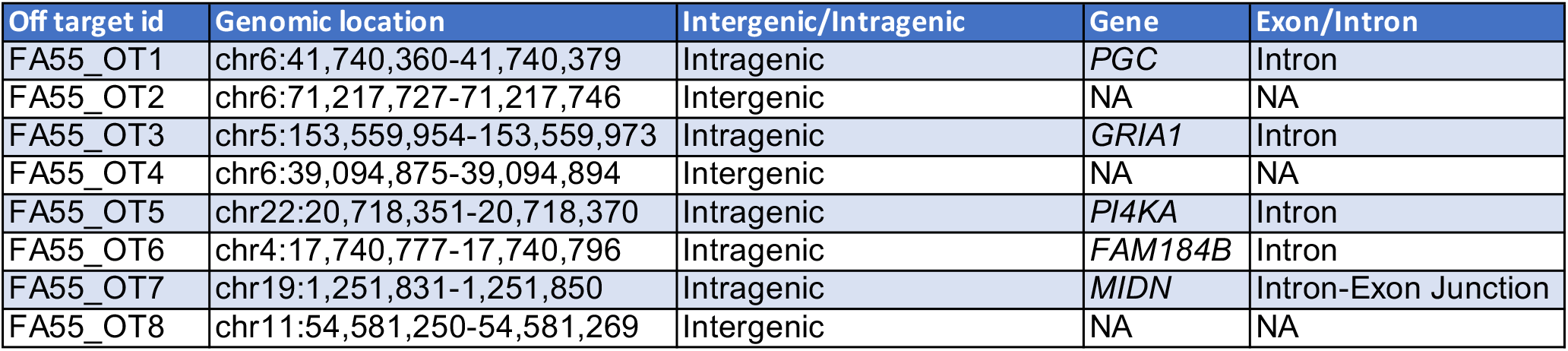
List of off-target sites for FA-55 sgRNA1 analysed in the study

| Off target id | Genomic location | Intergenic/Intragenic | Gene | Exon/Intron |
| --- | --- | --- | --- | --- |
| FA55_OT1 | chr6:41,740,360-41,740,379 | Intragenic | *PGC* | Intron |
| FA55_OT2 | chr6:71,217,727-71,217,746 | Intergenic | NA | NA |
| FA55_OT3 | chr5:153,559,954-153,559,973 | Intragenic | *GRIA1* | Intron |
| FA55_OT4 | chr6:39,094,875-39,094,894 | Intergenic | NA | NA |
| FA55_OT5 | chr22:20,718,351-20,718,370 | Intragenic | *PI4KA* | Intron |
| FA55_OT6 | chr4:17,740,777-17,740,796 | Intragenic | *FAM184B* | Intron |
| FA55_OT7 | chr19:1,251,831-1,251,850 | Intragenic | *MIDN* | Intron-Exon Junction |
| FA55_OT8 | chr11:54,581,250-54,581,269 | Intergenic | NA | NA |

**Table2:** List of off-target sites for FA75 sgRNA4 analysed in the study

| Off target id | Genomic location | Intergenic/Intragenic | Gene | Exon/Intron |
| --- | --- | --- | --- | --- |
| FA75_OT1 | chr2:110,027,307-110,027,506 | Intergenic | NA | NA |
| FA75_OT2 | chr2:110,344,019-110,344,038 | Intergenic | NA | NA |
| FA75_OT3 | chr2:202,148,657-202,148,676 | Intragenic | *KIAA2012* | Intron |
| FA75_OT4 | chr10:14,263,604-14,263,623 | Intragenic | *FRMD4A* | Intron |
| FA75_OT5 | chr9:42,184,196-42,184,215 | Intragenic | *SPATA31A6* | Intron |
| FA75_OT6 | chr14: 90,091,562-90,091,581 | Intragenic | *KCNK13* | Intron |
| FA75_OT7 | chr18:48,679,450-48,679,469 | Intragenic | *CTIF* | Intron |
| FA75_OT8 | chr3:63,777,216-63,777,235 | Intergenic | NA | NA |

**Table2:**
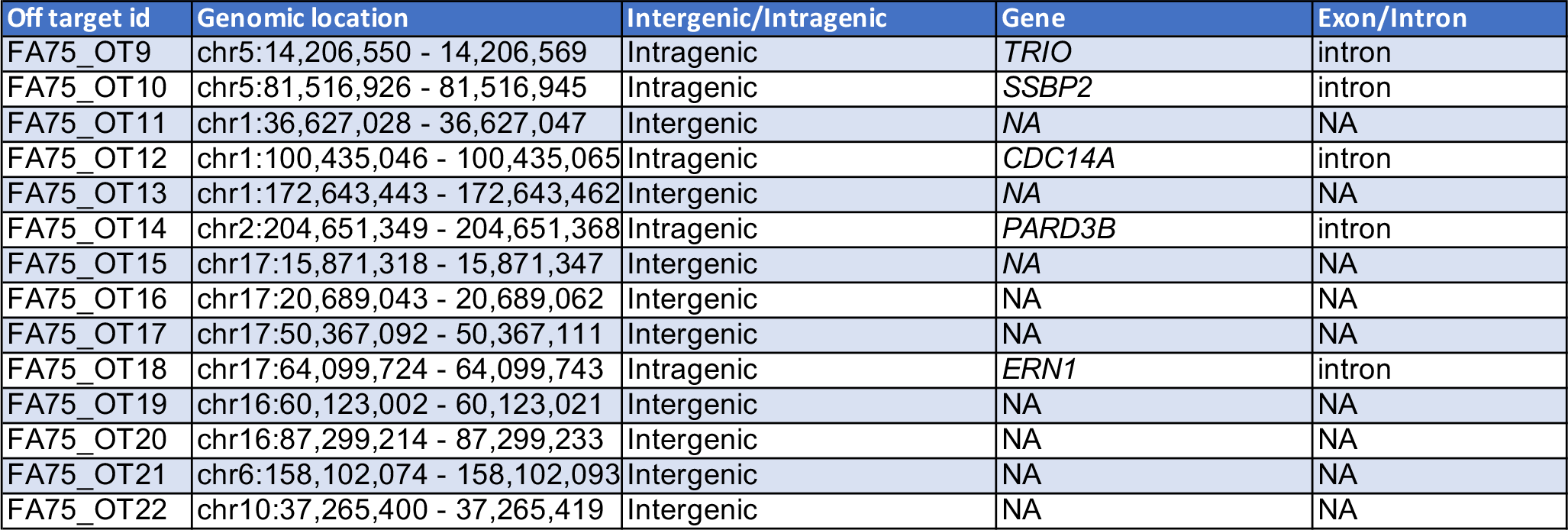
List of off-target sites for FA75 sgRNA4 not analysed in the study

**Figure 4:**
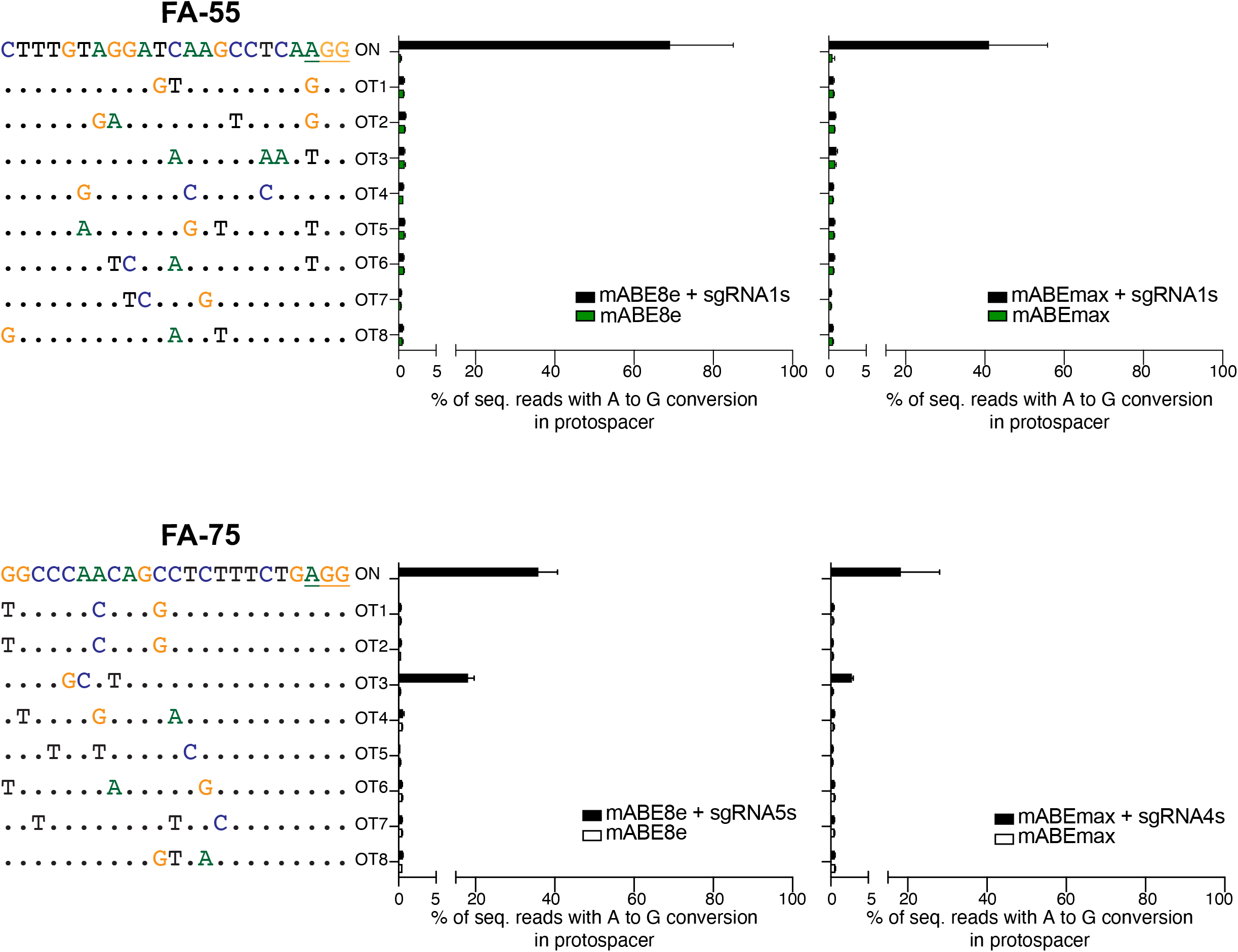
Off-target analysis at predicted locus in FA-55 and FA-75 LCLs. Computationally predicted off-target sites are shown for FA-55 sgRNA1 and FA-75 sgRNA4. Dots represent base identical to the on-target locus. PAM sites of FA-55 sgRNA1 and FA-75 sgRNA4 are underlined. Bars represent mean of amplicon NGS editing levels from 3 biological replicates, error bars indicate SD.

Given the promising results obtained in immortalized patient cells, we asked whether base editing approaches would potentially be suitable for preclinical models. Before moving to precious HSPCs from FA patients, we optimized electroporation conditions by targeting the *AAVS1* safe harbor locus with both ABEmax and ABE8e. We electroporated varying amounts of CD34^+^ cells from cord blood (CB) and mobilized peripheral blood (mPB) with mRNA forms of each base editor and a synthetic guide RNA targeting *AAVS1* (Figure 5A and Supp Figure A) ^40^. As in LCLs, ABE8e was much more efficient than ABEmax in both CB CD34^+^ cells (85±1.7% ABE8e vs 30.36±5.9% ABEmax) and the more clinically relevant mPB CD34^+^ cells (71.1±13.3% ABE8e vs 28±5.9% ABEmax) (Figure 5B and 5C).

**Figure 5:**
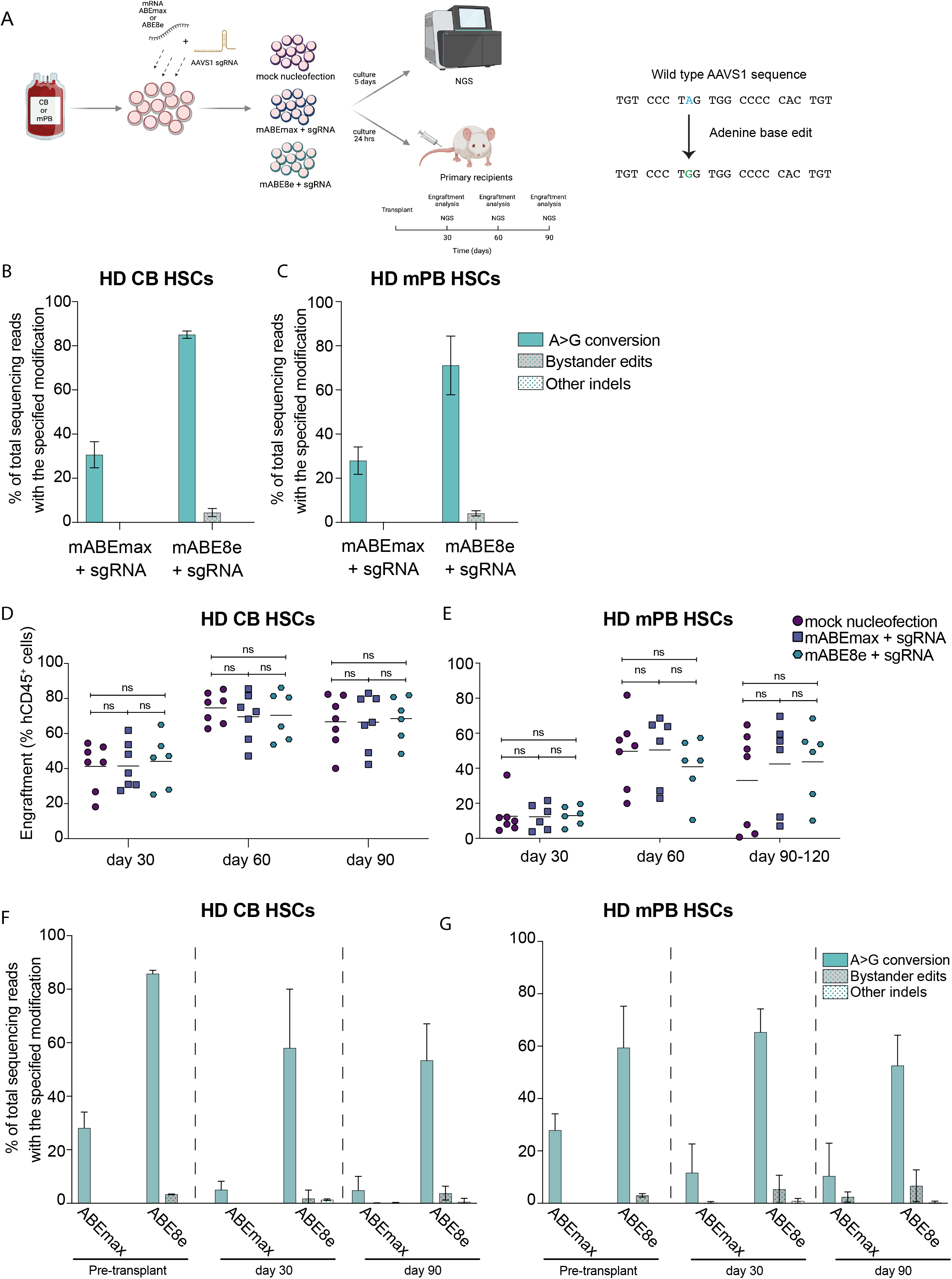
Adenine base edited human CD34+ cells successfully engraft into NSG mice. **A)** Schematics of experimental design to edit CD34^+^ primary cells. Healthy donor CD34+ cells from cord blood (CB) or mobilized peripheral blood (mPB) were purified by immunoselection and pre-stimulated 24 hours prior to base editing with mABEmax or mABE8e mRNA in combination with an *AAVS1* targeting synthetic gRNA. Edited CD34+ were maintained in culture 24 hours then transplanted into immunodeficient NSG mice. Pre-transplant amplicon NGS analysis was conducted 5 days after electroporation. **B)** Base editing frequencies at *AAVS1* in CB CD34^+^ cells edited with mABEmax or mABE8e measured by amplicon NGS. Bars represent mean of value from 3 biological replicates. **C)** Base editing frequencies at *AAVS1* in mPB CD34^+^ cells edited with mABEmax or mABE8e measured by amplicon NGS. Bars represent mean of value from 3 biological replicates. **D)** Human bone marrow engraftment and *in vivo* differentiation capacity of edited CD34^+^ cells from CB at 30, 60 and 90 days after transplant. Mean values are represented with a horizontal bar. In all cases, a two-way ANOVA was performed followed by a Tukey’s multiple comparison test: ns = not significant. **E)** Human bone marrow engraftment and *in vivo* differentiation capacity of edited CD34^+^ cells from mPB 30, 60 and 90or 120 days after transplant. Mean values are represented with a horizontal bar. In all cases, a two-way ANOVA was performed followed by a Tukey’s multiple comparison test: ns = not significant. **F)** Amplicon NGS analysis of editing levels in CD34+ CB cells (pre-transplant at 5 days post electroporation) and 30, 60 and 90 days post transplantation. Bars represent mean of value from 2 independent experiments with 4 to 7 mice per group, error bars represent SD. Pre-transplant data correspond to cells shown in Figure 5B. **G)** Amplicon NGS analysis of editing levels in CD34+ mPB cells (pre-transplant at 5 days post electroporation) and 30, 60 and 90 or 120 days post transplantation. Bars mean of value from 2 independent experiments with 4 to 7 mice per group, error bars represent SD. Amplicon NGS was conducted in 4 mice at 90 days post-transplant and 2 or 3 mice at 120 days post-transplant, per group. Pre-transplant data correspond to data shown in Figure 5B.

Analysis of individual hematopoietic colonies showed that ABE8e generated point conversions in homozygosis in all cases, confirming its efficiency in HSPCs from CB and mPB HD cells (Supp Figure D **and** E). However, NGS in liquid culture also revealed 4.2±1.3% bystander mutations in the targeted locus when using ABE8e. Regardless of the base editor, electroporation of base editor with synthetic guides into purified CD34^+^ cells did not cause gross defects in the clonogenic and differentiation potential of the HSPCs, suggesting that base editing was well tolerated in these cells ^41^(Figure 5 D **and** E **and** Supp Figure F **and** G).

To confirm that base editing can efficiently target long-term repopulating HSCs and does not affect the engraftment capacity of these precursors, unedited and edited CD34+ cells from CB and mPB sources from HDs were xenotransplanted into NOD.Cg-*Prkdc^scid^ Il2rg^tm1Wjl^*/SzJ (NSG) immunodeficient mice (Figure 5A). Monthly post-infusion, BM cells were collected from transplanted recipients by femoral BM aspiration. Human engraftment was analyzed by flow cytometry using anti-hCD45-FITC to detect human engraftment. Multilineage reconstitution was assessed using anti-hCD34-APC, anti-hCD33-PE, anti-hCD19-Pe-Cy5, and anti-hCD3-Pe-Cy7.

We found that CB HSPCs edited with either ABEmax or ABE8e engrafted with similar efficiencies to mock-edited counterparts (median levels of engraftment of 66.8±15.2% mock; 66.5±15.9% ABEmax; 68.5±13.0% ABE8e) at three months post-infusion (Figure 5D). As expected, engraftment of mPB CD34^+^ cells was lower than for CB, but also comparable between mock and base edited cells (33.1±28.1 mock; 42.4±26.2 ABEmax; 43.7±21.7 ABE8e) (Figure 5E). No overt toxicity associated to the treatment was observed in these mice (Figure 5D and E). Also the proportion of myeloid and lymphoid lineages and hCD34^+^ cells repopulating the BM of transplanted recipients were similarly represented among unedited and the two type of edited cells (Figure 5D and 5E, Supp Figure G **and** H).

Deep sequencing analysis after long-term engraftment of edited hCD34^+^ cells showed that ABE8e more efficiently targeted HSCs in comparison to ABEmax, reaching median values of editing higher than 50% at 3-4 months post-transplant, regardless of the HSC source, CB or mPB (Figure 5F and 5G).

Finally, we investigated whether base editing approaches could correct mutations in HSPCs from FA patients. Because of the extreme scarcity of HSPCs in FA patients, we tested the efficiency of gene editing in Lineage depleted (Lin-) cells from a patient carrying the *FANCA* c.295C>T (FA-55) mutation (Figure 6A). Since ABE8e editing of FA-55 exhibited high on-target activity, no bystander modifications and no off-target activity in LCLs (Figure 3B), this base editor was selected to target Lin-cells from this patient.

**Figure 6:**
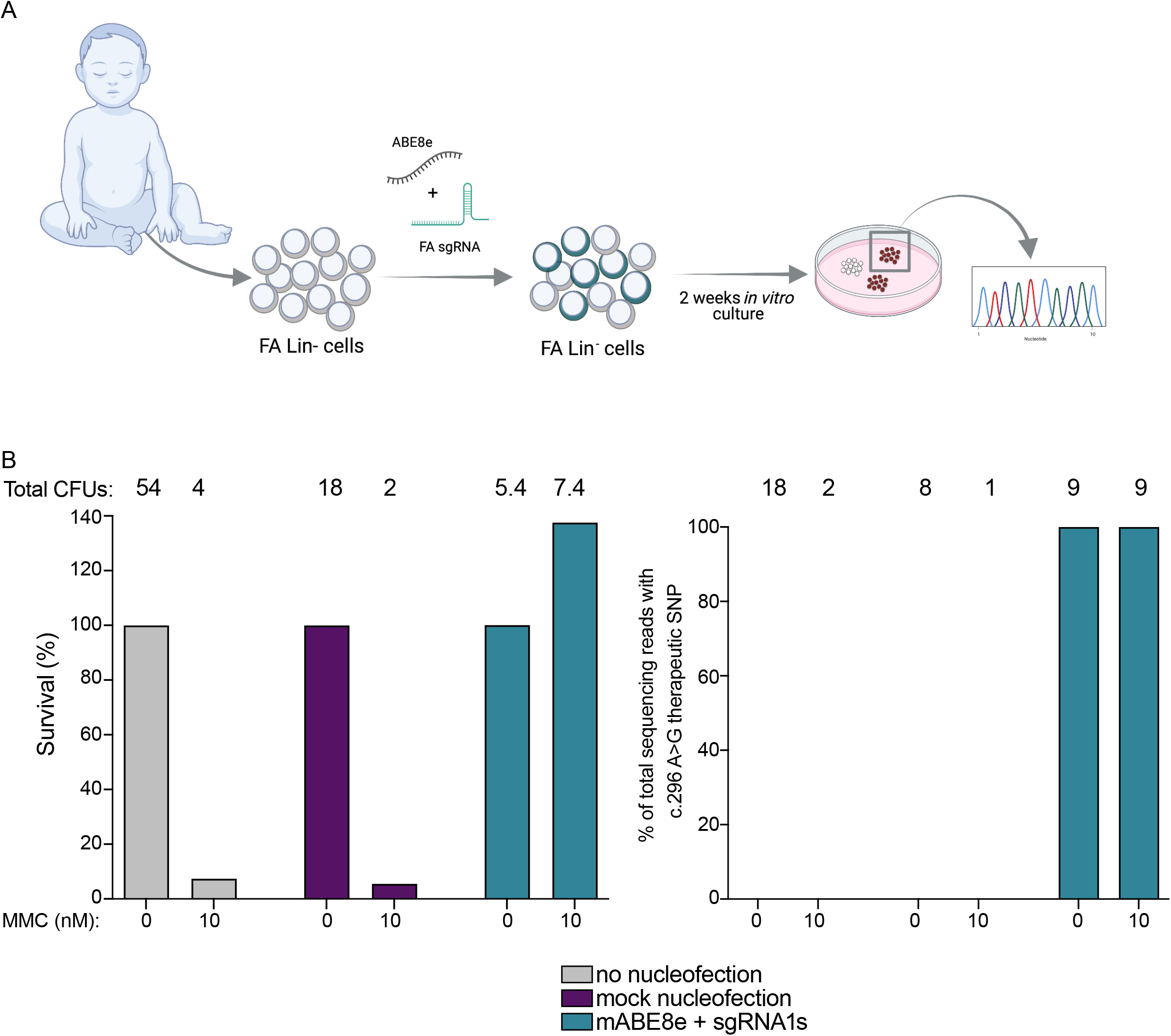
mPB lineage negative cells from a FA-A patient (FA-55) can be efficiently edited by ABE8e. **A)** Schematic representation of the experiment to edit lineage depleted (lin-) cells from an FA patient harboring c.295 C>T mutation. FA Lin-HSPCs were pre-stimulated for 24hr and edited with mABE8e mRNA and synthetic FA-55 sgRNA1. 24 hours after electroporation, CFUs were plated and number of CFUs in the absence or presence of MMC were scored. Individual colonies were collected to analyze the percentage of edited CFUs. **B)** Survival of hematopoietic colonies (CFUs) obtained from unedited (no electroporation or mock electroporation) and edited FA Lin-HSPCs before and after MMC exposure (10nM). Number of CFUs in each condition is indicated above bars. **C)** Sanger sequencing analysis of individual colonies from patient FA-55 after base editing with ABE8e in the absence or presence of MMC.

Thawed Lin-cells were pre-stimulated for 24 hours and electroporated with ABE8e mRNA together with a synthetic FA-A targeting gRNA. At 24 hours after electroporation, we seeded untreated, mock nucleofected and mABE8e edited cells in methylcellulose and treated each culture with low dose (10 nM) MMC or vehicle. In the presence of MMC, mock electroporated or non-electroporated Lin-cells exhibited very poor colony forming ability (Figure 6B). By contrast, ABE8e edited Lin-cells generated similar number of colonies in the presence and absence of MMC, confirming the reversion of the characteristic MMC hypersensitivity of FA cells (Figure 6B). Sanger sequencing in individual CFCs showed that gene editing occurred at both alleles in all the colonies analyzed, regardless that these colonies were generated in the presence of the absence of MMC (Figure 6C). These data demonstrate the high efficiency of ABE8e to target HSPCs from HDs and FA patients, and highlights the potential of base editors to correct a prevalent mutation observed in FA.

## Discussion

Here we explored the possibility of using CRISPR-Cas base editors to reverse the effects of FA mutations in patient-derived LCLs and HSPCs. While NHEJ-and HDR-based strategies have been explored to genetically treat FA HSPCs ^18, 42–44^, this is the first study to our knowledge that demonstrates proof-of-concept that base editors are tolerated and highly efficient in FA HSPCs.

The FA pathway is multi-functional, with roles in DNA crosslink repair, DSB repair and replication fork restart, among other relevant functions ^45–47^. Lack of these activities has compromised HDR-based approaches for allele correction in FA patient cells. However, we found that absence of functional FA pathway did not interfere with adenine base editor activity in LCLs and HSPCs. This suggests that the FA pathway is dispensable for base editor activity and could be exploited as a novel therapeutic strategy in FA. Both *FANCA* nonsense-to-missense and missense-to-wildtype editing resulted in phenotypic rescue on both the molecular and phenotypic level. We are optimistic that the approach outlined here could be extended to additional FA alleles to form the foundation of future gene editing therapies for FA.

We found that ABE editing can yield phenotypic correction in FA LCLs and FA HSPCs. In edited FA-55 LCLs, we observed re-expression of FANCA protein expression, molecular evidence of FA pathway re-activation, and a significant proliferative advantage over unedited cells. Similarly, edited FA-75 LCLs showed phenotypic correction on multiple levels. Base edited FA HSPCs grow in the presence of MMC, indicating phenotypic correction of primary patient cells. However, access to sufficient numbers of FA HSPCs prevented us from currently determining whether these phenotypically corrected FA HSPCs efficiently engraft in an immunodeficient mouse model. Such a test will be an important step in further preclinical studies. Importantly, engraftment capacity of base edited HSPCs in immunodeficient mice was not altered in comparison with uneditied cells either with from HD CB and mPB CD34+ cells were used. These results are consistent with recent studies performed in healthy and sickle cell HSPCs ^41, 48, 49^. Further studies will directly address the important question of engraftment potential for base edited FA cells.

The extremely high activity of ABE8e might result in higher levels of unintended mutations at the editing window and at off-target loci ^35^. We detected high levels of unintended mutations in the FA-75 editing window, but not at the FA-55 site. We also found a prominent ABE8e off-target at a candidate site for the FA-75 sgRNA, but none for the FA-55 sgRNA. Despite unintended modifications in ABE8e edited cells, it was capable of much higher editing efficiency in CD34^+^ HSPCs (Figure 5), thus enabling editing of clinically-relevant long term HSPCs. ABE8e could still be valuable in FA-75, since the bystander edits occur at wobble bases and are predicted to be neutral once the corrected allele expands. Our phenotypic analysis in FA-75 LCLs (Figure 3) indicated that these bystander mutations do not affect FANCA function and FA pathway activation. If needed, one could reduce the bystander and off-target actives of ABE8e by using an ABE8e RNP or ABE8e virus-like particles, which have been reported to reduce off-target DNA effects ^35, 50^. The ABE8e (TadA-8e V106W) variant could also be a useful tool to reduce unintended base modification. Unbiased identification of genomic and transcriptomic off-targets will be an important next preclinical step in validating any base editing approach to cure FA.

Overall, our study indicates that adenine base editing is a feasible approach for the efficient restoration of function in FA patient HSPCs. These results provide a foundation for the use of base editors in FA and other DNA repair disorders, where these targeted tools may be both more efficacious and even safer, as compared to current untargeted gene addition therapies.

## Materials and Methods

### Plasmid generation

sgRNAs were designed to contain a variable 20 nucleotide sequence, corresponding to the target gene. Oligos for sgRNAs and nicking guides were ordered from IDT and cloned into the pLG1-puro-BFP vector after digestion with BstXI and BlpI. Base editor plasmid ABEmax_P2A_GFP (Addgene plasmid # 112101) was a gift from David Liu and Lukas Villiger. The coding sequences of ABEmax or ABE8e were cloned into a T3 promotor containing pRN3 plasmid using the NEBuilder® HiFi DNA Assembly Master Mix (New England Biolabs). All vectors were purified using Qiagen Spin Mini- or Midiprep (Qiagen) with endotoxin removal step. Primers used in this study can be found in the oligo table.

### mRNA production for ABE base editors

All mRNA used in this study was generated by the following synthesis protocol. The mRNA template plasmid was linearized by digestion with SfiI (50°C, overnight) and 200 µl of the digestion reaction was combined and mixed 1:1 with phenol chloroform for extraction. Samples were vortexed for 15 sec. at high speed and then centrifuged at 13,000×g for 5 min. 150 µl of the aqueous phase were transferred into a new tube and 1:10 volume of 3 M NaOAc and 165 µl of isopropanol were added. After 30 min incubation at −80°C the samples were centrifuged at 4°C, top speed for 30 min. The supernatant was carefully removed while not disturbing the pellet and 400 µl of 80% EtOH were added for another spin of 5 min. The EtOH was removed without leaving residuals and the pellet was dissolved in 10 µl of RNAse-free water. For in vitro transcription, the mMESSAGE mMACHINE® T3 Kit (Life technologies) was used as described in the user guide. 1 µg of linear plasmid was used as a template and transcription reaction was carried out for 2 h at 37°C. For removal of the residual DNA template, 1 µl of TURBO DNAse was added to the transcription reaction for 15 min. RNeasy Mini Kit (Qiagen) was used for cleanup of the transcription reaction. In vitro transcribed mRNAs were kept at −80°C until further use.

### Cell lines

Patient-derived LCLs (FA-55 and FA-75) and HD-LCLs were a gift from Dr. Paula Rio, (CIEMAT, Spain). LCLs were cultured in Roswell Park Memorial Institute medium (RPMI from ThermoFisher Scientific) supplemented with 20% Hyclone fetal bovine serum (FBS), 1% penicillin/streptomycin (P/S) solution, 0.005mM β-mercaptoethanol and 1% non-essential amino acids. Cells were split every two days to keep them at a density of 5×10^5^ cells/ml in 37°C, 5% CO_2_.

### Editing LCLs with base editor plasmids

For base editing experiments, LCLs were run through Ficoll gradient and the death cells and debris were cleared. 5×10^5^ LCLs were electroporated with ABEmax (750 ng) and sgRNA (250 ng) using 4D-Nucleofector™ X unit from Lonza (SF solution, DN100 (FA-55) and CM137 (FA-75)). Cells were cultured in a 24 well dish after nucleofection and transferred into a T25 flask after recovery for the long term culturing.

### Editing LCLs with in vitro transcribed mRNA

For base editing with ABEmax mRNA, 2×10^5^ FA-55 or FA-75 LCLs were electroporated with 3 µg or 6 µg BE mRNA and 100 or 200 pmol of synthetic sgRNAs (Synthego), respectively. For both experiments the Lonza nucleofector was used with SF solution and the EW113 nucleofection program. Nucleofection efficiency and cell viability were assessed by flow cytometry 24 hours after the nucleofection. Cells were cultured in a 96 well dish after nucleofection and transferred into a T25 flask later.

### Sanger and Next generation (NGS) sequencing

Genomic DNA was extracted using QuickExtract™ DNA Extraction Solution (Lucigen) and genomic locus of the interest was amplified by using AmpliTaq Gold^®^ 360 Master Mix (ThermoFisher Scientific). Primers for PCR and Sanger sequencing can be found in the supplemental table 1. For NGS library preparation two rounds of PCR were performed. In the first one (PCR 1), the PCR primers contained the corresponding sequence to the genomic locus and the appropriate forward and reverse Illumnia adapters sequences (supplemental table 1). PCR 2 was carried out with unique Illumina barcoding primer combinations using 15 µl of purified product from PCR1. PCR2 was purified by SPRIselect beads (Beckman). A ratio of 0.9× beads/PCR product volume was used. The resulting amplicon size and concentration was verified on the 4200 TapeStation System (Agilent) before multiplexing. For Sanger sequencing (and PCR1 for NGS) the products of the PCR were purified using MinElute columns (Qiagen) and eluted in 30 µl elution buffer (EB). ∼120 ng of purified PCR product was sent for Sanger economy sequencing. The forward primer was used for sequencing FA-55, while the reverse one was used for FA-75. Sanger sequencing graphs were generated using Geneious Prime 2020.2.3.

### Western blotting and MMC treatment

For MMC treatment, 2×10^6^ LCLs were incubated with 1 µM MMC for 24 h before 1×10^6^ cells were collected. For protein extracts, 1×10^6^ LCLs were pelleted and washed in PBS. To lyse the cells, 150 µl of ice-cold RIPA buffer (Millipore) supplemented with Halt protease inhibitor cocktail (ThermoFisher Scientific) was used. LCLs were resuspended in this lysis buffer and incubated on ice for 20 min. After centrifugation at 22,000 rpm for 30 minutes at 4 °C the supernatant was transferred into another microcentrifuge tube. Protein concentration was measured, using Bradford Assay (VWR) and after incubation in RIPA supplemented with 1×LDS and 1×DTT for 5 minutes at 95 °C, 15 µg protein were loaded on the gel. Gel electrophoresis was run with 4-12% polyacrylamide gels (NuPAGE) and 1×MOPS SDS running buffer (NuPAGE). Proteins were transferred using the Criterion Trans-Blot^®^ Cell (BioRad) with a Tris-Glycine transfer buffer (25 mM Tris base, 192 mM glycine, 20% methanol (v/v); pH = 8.3). Membrane was incubated with Ponceau staining for a few minutes to confirm transfer and then blocked with 5% (w/v) non-fat dry milk in TBS-T (0.1% Tween-20) for 1 hour at room temperature. Primary anti-rabbit FANCA antibody (ab5036, Abcam or bethyl), anti-rabbit FANCD2 antibody (ab221932, Abcam), anti-goat HSP60 antibody (sc-1052, Santa Cruz Biotechnology) were diluted 1:1,000 in 10% milk TBS-T. HSP60 served as a loading control. Membrane was stained with antibody overnight at 4°C and then washed three times in TBS-T before 45 min incubation with anti-rabbit secondary antibody (IRDye 800CW (926-32213) or anti-goat IRDye 800CW (926-32214), 1:5,000 diluted in 10% milk TBS-T. Finally, the membrane was washed two times with TBS-T and one time with PBS before imaging with the Li-Cor’s Near-InfraRed fluorescence Odyssey CLx Imaging System.

### MMC sensitivity assay

MMC sensitivity assay was performed, incubating 2.5×10^5^ cells for 5 days in media with increasing concentrations of MMC (0, 3, 10, 33, 100, 333, 1000 nM). Survival was measured by flow cytometry using the forward and side scatter to gate for the life cell population. Downstream analysis was performed using FlowJo Software v10.7.1 (FlowJo, LLC). Each data point represents the mean of three biological replicates.

### NGS data analysis

Demultiplexing of the Sequencing reads was done with the MiSeq Reporter (Illumina). Sequencing reads were aligned to the genome using the bowtie2 algorithm and visualized using the Integrative genome viewer. CRISPResso2 was run in with the following settings: CRISPRessoBatch --batch_settings batch.batch --amplicon_seq -p 4 --base_edit -g -wc -10 -w 20. Corrected reads with the base edited therapeutic SNP were calculated by selecting only reads with the intended edit but no indels in the quantification window. Percentages of corrected read and uncorrected reads were plotted using GraphPad Prism 8.3.1 (GraphPad Software, Inc., San Diego, CA).

### Off-target analysis for base editing FA-55 with ABEmax mRNA

Cas-OFFinder (http://www.rgenome.net/cas-offinder/) ^39^ was used to determine all possible off target sites. Cas-OFFinder was run under the following settings: mismatch number = 3 (equal or less), DNA Bulge Size = 0 and RNA Bulge Size = 0. NGS was performed on the respective genomic sites using NGS primers listed in Sup. Table 1. Data were analyzed by CRISPResso2 and run with the same setting as for on target base editing. For quantification of A to G conversions, all adenines in the protospacer were considered potential targets of the BE. Therefore, all reads which contained one or more A to G conversion in this window were scored as base edited and the sum of all reads with A to G conversions at these positions was calculated.

### Protein sequence alignment

Protein sequences for FANCA were retrieved from https://w.ncbi.nl.nih.gov/protein and converged together. A multiple sequence alignment was created using T-Coffee (http://tcoffee.crg.cat/apps/tcoffee/do:regular) and was visualized with the help of Boxshades (https://embnet.vital-it.ch/software/BOX_form.html). Using the “fasta_aln” result file from T-Coffee with format “other” as input and “RTF_new” as the output format.

### Hematopoietic stem and progenitor cells from healthy donors and FA patients

Human CD34^+^ cells were obtained from healthy donor umbilical cord blood (UCB) or mobilized peripheral blood samples provided by *Centro de Transfusiones de la Comunidad de Madrid* and *Hospital Niño Jesús*, respectively. Mononuclear cell fractions were purified by Ficoll-Paque PLUS (GE Healthcare) density gradient centrifugation according to manufacturer’s instructions. Human CD34^+^ HSPCs were purified from the mononuclear fraction by immunoselection using the CD34 Micro-Bead Kit (MACS, Miltenyi Biotec). Magnetic-labelled cells were selected with a LS colum in *QuadroMACS^TM^ Separator* (Miltenyi Biotec) following manufacturer’s instructions. Purified hCD34^+^ were then analysed by flow cytometry to evaluate their purity in *LSRFortessa Cell Analyser* (BD) using *FlowJo Software* v10.7.1. Purities ranging from 85-98% were routinely obtained. Cells were grown in *StemSpam* (StemCell Technologies) supplemented with 1% GlutaMAX™ (Gibco), 1% P/S solution (Gibco), 100 ng/mL human stem cell factor (hSCF), human FMS-like tyrosine kinase 3 ligand (hFlt3-L), human thrombopoietin (hTPO), and 20 ng/mL human interleukin 3 (hIL3) (all obtained from EuroBiosciences) under normoxic conditions. HSPCs were pre-stimulated 24 hours prior electroporation. Cryopreserved CD34+ cells were thawed and cultured under the same conditions 24 hours prior electroporation.

Lineage negative populations from FA patients were obtained from apheresis aliquots by the incubation of cells with anti-hCD3-PE, anti-hCD19-PE, anti-hCD33-PE, antih-CD-235a PE for 30 min. Then cells were washed and incubated with anti-PE Microbeads (Miltenyi Biotec). Lineage negative population was confirmed in *LSRFortessa Cell Analyser* (BD) using *FlowJo Software* v10.7.1. Cells were grown and cultured during 24 hours prior electroporation in GMP Stem Cell Grow Medium (CellGenix) supplemented with 1% GlutaMAX™ (Gibco), 1% P/S (Gibco), 100 ng/mL SCF and Flt3, 20 ng/mL TPO and IL3 (EuroBiosciences), 10 µg/mL anti-TNFα (Enbrel-Etanercept, Pfizer) and 1 mM N-acetylcysteine (Pharmazam) under hypoxic conditions (37°C, 5% of O_2_, 5% of CO_2_ and 95% RH).

### mRNA electroporation

Electroporation was performed using Lonza 4D Nucleofector (V4XP-3032 for 20-μl Nucleocuvette Strips or V4XP-3024 for 100-μl Nucleocuvette Strips) according to the manufacturer’s instructions. The modified synthetic sgRNA (2-O-methyl 3′ phosphorothioate modifications in the first and last three nucleotides) were purchased from Synthego and BE mRNA was obtained through in vitro transcription using mMESSAGE mMACHINE^TM^ T3 Transcription kit (Invitrogen). 2×10^5^ HSPCs from healthy donor were resuspended in 20 µL P3 solution and electroporated in 20-µL Nucleocuvette wells using program EO-100 with increasing concentration of BE mRNA and sgRNA (3 µg of BE mRNA and 3.2 µg sgRNA for HD CB cells and 6 µg of BE mRNA and 6.4µg sgRNA for HD mPB cells). For 100-µL cuvette electroporation, 1×10^6^ HSPCs were resuspended in 100 µL P3 solution and electroporated using 30 µg of BE mRNA and 32 µg of sgRNA with program EO-100. FA Lineage negative cells were electroporated using similar conditions. Electroporated cells were resuspended in *StemSpam* medium (StemCell Technologies) with corresponding cytokines. Then, 24 hours later, cells were used for transplant or maintained in culture for 5 days for DNA extraction and sanger/NGS analysis to evaluate basal gene editing.

### Colony Forming Unit Assay

Colony forming unit assays were established using 900 HD hCD34^+^ or 7.4×10^4^ FA-A hLin^+-^ cells in 3 mL of enriched methylcellulose medium (StemMACS™ HSC-CFU complete with Epo, Miltenyi Biotech). In the case of FA cells, 10 μg/mL anti-TNFα and 1mM N-acetylcysteine were added. Each mL of the triplicate was seeded in a M35 plate and incubated under normoxic (HD hCD34^+^ cells) or under hypoxic (FA hLin^-^ cells) conditions. To test MMC sensitivity of hematopoietic progenitors obtained from FA-A patients, 10nM of MMC (Sigma-Aldrich) was added to the culture. After fourteen days, colonies were counted using an inverted microscope (Nikon Diaphot, objective 4×) and CFUs-GMs (granulocyte-macrophage colonies) and BFU-Es (erythroid colonies) were identified.

### Base editing efficiency measurement in HSPCs by NGS

Base editing frequencies were measured either from liquid cultures 5 days after electroporation or in individual hematopoietic colonies grown in methylcellulose. The *AAVS1* or *FANCA* exon 4 regions were amplified with AmpliTaq Gold 360 DNA Polymerase (Thermo Fisher Scientific) and corresponding primers using the following cycling conditions: 95°C for 10 min; 40 cycles of 95°C for 30s, 60°C for 30s and 72°C for 1min; and 72°C for 7min. Primers used in these PCRs are listed in Table 1. Resulting PCR products were subjected to sanger sequencing or illumina deep sequencing. For Sanger sequencing, PCR products were sequenced using Fw primers described in Table 1. For deep sequencing, PCR products were purified using the Zymo Research DNA Clean and Concentrator kit (#D4004), quantified using Qubit fluorometer (Thermo Fisher Scientific), and used for library construction for illumine platforms. The generated DNA fragments were sequenced by Genewiz with *Illumina MiSeq Platform*, using 250-bp paired-end sequencing reads. Frequencies of editing outcomes were quantified using CRISPResso2 software (quantification window center (−3) and size (−10); plot window size (20); base edit target A to G; batch mode).

### Base edited HSPCs transplantation studies in NSG mice

HD hCD34^+^ cells from CB or mPB were purified and pre-stimulated for 24 hours for electroporation as previously described. Three groups of cells were established: electroporated cells without nuclease or sgRNA (Mock); electroporated cells with ABEmax mRNA and sgRNA (ABEmax); and electroporated cells with ABE8a mRNA and sgRNA (ABE8a). Twenty-four hours later, 3×10^5^ cells per mouse were intravenously injected into immunodeficient NSG mice previously irradiated with 1.5 Gy. A CFU-assay was also conducted and the remaining cells were pelleted for DNA extraction and NGS analysis to evaluate basal gene editing. 30 and 60 days after transplantation, bone marrow samples were obtained by intra-femoral aspiration and total human engraftment was measured by flow cytometry, analysing percentage of hCD45^+^ cells (anti-hCD45-FITC, BioLengend). Multilineage reconstitution was also evaluated using antibodies against hCD34 (anti-hCD34-APC, BD) for HSPCs, hCD33 (anti-hCD33-PE, eBioscience) for myeloid cells, hCD19 (anti-hCD19-Pe-Cy5, BioLegend) for B cells and hCD3 (anti-hCD3-Pe-Cy7, BioLegend) for T cells. The remaining cells were pelleted for DNA extraction and NGS analysis to evaluate the presence of gene edited cells. Mice were euthanized at 90 or 120 days post-transplantation, and bone marrow cells were obtained from hind legs. Human engraftment was evaluated by flow cytometry according to the percentage of hCD45^+^ cells in the different hematopoietic organs. Multilineage reconstitution was determined using antibodies against hCD34 for HSPCs, hCD33 for myeloid cells, hCD19 for B cells and hCD3 for T cells. Viable cells were identified by 4’, 6-diamidino-2-phenylindole (DAPI). Flow cytometry analysis were performed using a LSRFortessa Cell Analyzer and analyzed with FlowJo Software v10.7.1.

---

**Supplemental Figure to figure 5:**
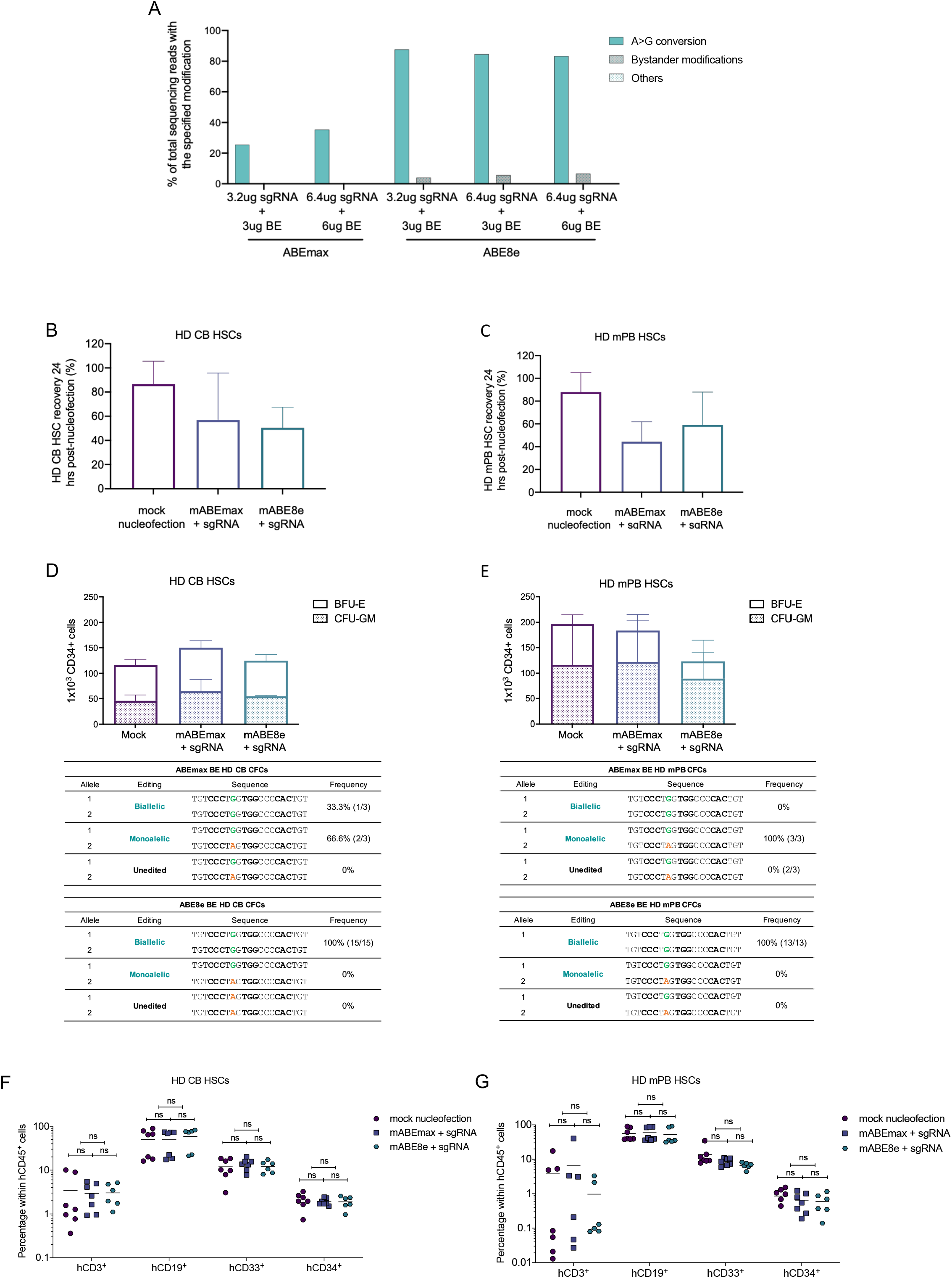
Adenine base editing efficiency in healthy donor (HD) CD34+ cells. **A)** Optimization of gene editing efficiencies using mABEmax and mABE8e base editors in HD CB CD34+ cells. **B and C)** CB and mPB cell recovery analyzed 24 hours after mock, mABEmax and mABE8e mRNA and synthetic gRNA electroporation. Bars represent mean value from two independent experiments, error bars indicate SD. Data correspond to cells shown in Figure 5B and C. **D and E)** Upper: number of hematopoietic myeloid and erythroid colonies (CFU-GM and BFU-E, respectively) per 1×10^3^ HD CD34^+^ cells after mock, mABEmax and mABE8e mRNA and synthetic gRNA electroporation in CB and mPB CD34^+^ cells. Lower: Analysis of the frequency of specific editing events in human hematopoietic colonies (Total CFUs: CFU-GM + BFU-E). Bars represent mean value of three experimental replicated from one independent experiment (**D**) and two independent experiments (**E**), error bars indicate SD. Data correspond to cells shown in Figure 5B and C. **F and G**) Multilineage repopulation of human HSPCs (HD CB HSCs or HD mPB HSCs) in bone marrow from recipient mice 90-120 days post-transplant. Mean values are represented with a horizontal bar. In all cases, a two-way ANOVA was performed followed by a Tukey’s multiple comparison test: ns = not significant.

## Acknowledgements

JEC is a cofounder and board member of Spotlight Therapeutics, a co-founder and SAB member of Lyrik Therapeutics, an SAB member of Mission Therapeutics, an SAB member of Relation Therapeutics, an SAB member of Hornet Bio, an SAB member for the Joint AstraZeneca-CRUK Functional Genomics Centre, and a consultant for Cimeo Therapeutics. The lab of JEC has funded collaborations with Allogene. JEC is supported by the NOMIS Foundation and the Lotte and Adolf Hotz-Sprenger Stiftung. MEK is supported by the Fanconi Anemia Research Foundation.

We thank the Functional Genomics Center Zurich (FGCZ) and especially Dr. Susanne Kreutzer and Dr. Zacharias Kontarakis for their help for NGS sequencing. We thank Lukas Villiger for sharing ABEmax-GFP plasmid and we thank the members of the Corn Lab for helpful discussions and help with the manuscript.

## Author Contributions

SMS, MEK, PR, JEC conceived this project. SMS, AC performed the experiments from Figure 1 to Figure 4 LU performed the experiments Figure 5, 6 and supplemental figure with the help of LGG in mouse experiments. MEK, JEC wrote first manuscript with the contributions from PR and LU and other authors. All authors read and approved the final manuscript.

---

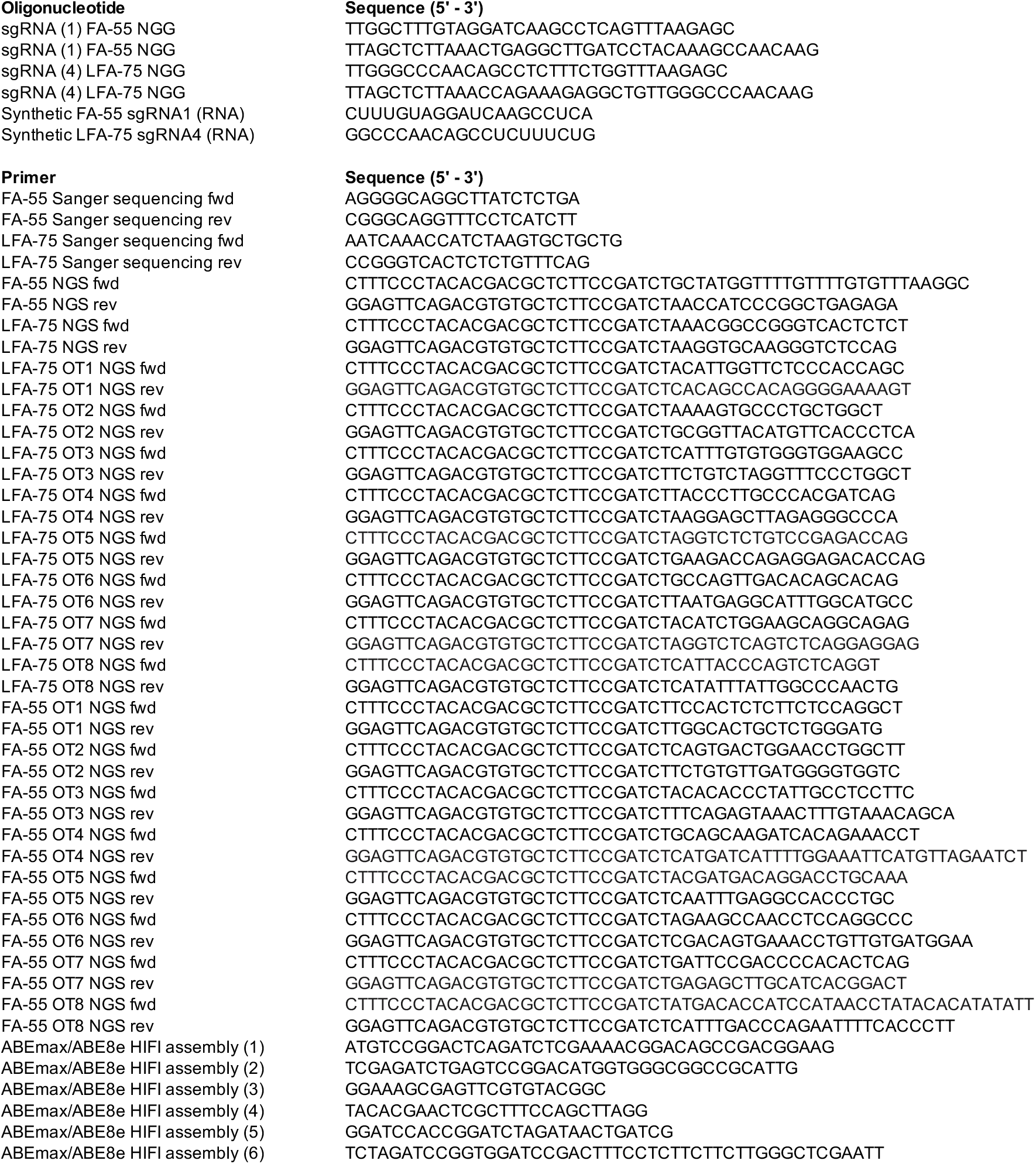

